# Connectome-based model predicts individual psychopathic traits in college students

**DOI:** 10.1101/2021.06.21.449277

**Authors:** Shuer Ye, Bing Zhu, Lei Zhao, Xuehong Tian, Qun Yang, Frank Krueger

**Author notes:** Corresponding authors **Qun Yang, Ph.D.** Institute of Psychological Sciences, Hangzhou Normal University, 2318 Yuhangtang Road, Hangzhou, China.

## Abstract

**Background:** Psychopathic traits have been suggested to increase the risk of violations of socio-moral norms. Previous studies revealed that abnormal neural signatures are associated with elevated psychopathic traits; however, whether the intrinsic network architecture can predict psychopathic traits at the individual level remains unclear.

**Methods:** The present study utilized connectome-based predictive modeling (CPM) to investigate whether whole-brain resting-state functional connectivity (RSFC) can predict psychopathic traits in the general population. RS functional magnetic resonance imaging data were collected from 84 college students with varying psychopathic traits measured by the Levenson Self-Report Psychopathy Scale (LSRP).

**Results:** We found that RSFC of the negative networks predicted individual differences in total LSRP and secondary psychopathy scores but not primary psychopathy score. Particularly, nodes with the most connections in the predictive connectome anchored in the prefrontal cortex (e.g., anterior prefrontal cortex and orbitofrontal cortex) and limbic system (e.g., anterior cingulate cortex and insula). In addition, the connections between the occipital network (OCCN) and cingulo-opercular network (CON) served as a significant predictive connectome for total LSRP and secondary psychopathy score.

**Conclusion:** CPM constituted by whole-brain RSFC significantly predicted psychopathic traits individually in the general population. The prefrontal cortex and limbic system at the anatomic level and the CON and OCCN at the functional network level plays a special role in the predictive model—reflecting atypical executive control and affective processing for individuals with elevated psychopathic traits. These findings may provide some implications for early detection and potential intervention of psychopathic tendency.

## Introduction

Psychopathy is a dimensional personality construct which is characterized by a discernible cluster of personality traits, including shallow affect, reduced control of impulse, high risk taking, pathological lying (Douglas, Nikolova, Kelley, & Edens, 2015). Individuals with elevated psychopathic traits demonstrate increased maladaptive behavior, such as violence and aggression, and therefore have negative impacts on the stability and security of society (R. D. Hare, 2006). Psychopathic traits are captured by two factors— primary and secondary psychopathy which are linked to different behavioral features and may be related to different causes (Lee & Salekin, 2010). Primary psychopathy is mainly associated with interpersonal and affective features which has been widely proved to be related to emotional deficits and are more likely to originate from neurobiological dysfunctions. In contrast, secondary psychopathy is more associated with the lifestyle/antisocial feature which is closely related to impulsivity and antisocial behavior and are more likely to root in environmental causes (Skeem, Johansson, Andershed, Kerr, & Louden, 2007; Yildirim & Derksen, 2015).

Despite that previous research has identified dysfunctional neurobiological signatures for people with elevated psychopathic traits (Cohn et al., 2015; Korponay et al., 2017), few has examined whether intrinsic network architecture can predict psychopathic traits at the individual level by using resting-state functional magnetic resonance imaging (rs-fMRI). Investigations on prediction about individualized psychopathic traits using rs-fMRI may increase our understanding of the neurobiological substrates associated with psychopathic traits and may provide some implications for early detection and potential intervention for psychopathic tendency in the future.

Resting-state functional connectivity (RSFC) —defined as temporal dependency of low-frequency (0.01–0.1 Hz) BOLD fluctuations between anatomically separated brain regions at the resting state— has the potential to serve as neural fingerprints of human behavior (Friston, Frith, Liddle, & Frackowiak, 1993; Van Den Heuvel & Pol, 2010). RSFC has been validated to be a good predictor of personality traits, social behaviors, and psychiatric disorders (Finn et al., 2015; Gordon et al., 2017). Psychopathic traits are related to decreased RSFC between brain regions involved in affective processing (Motzkin, Newman, Kiehl, & Koenigs, 2011), social cognition (Glenn & Raine, 2008), and moral decision making (Pujol et al., 2012). In particular, the prefrontal cortex (PFC) is typically associated with psychopathic traits (N. E. Anderson & Kiehl, 2012; Viding & McCrory, 2018). For example, the ventromedial prefrontal cortex (vmPFC), a brain region responsible for value evaluation, emotion regulation, and moral judgment, is found to be dysfunctional in individuals with elevated psychopathic traits (R. J. R. Blair, 2007; Gangopadhyay, Chawla, Dal Monte, & Chang, 2020; T. A. Hare, Camerer, & Rangel, 2009).

Further, RSFC in the dorsolateral prefrontal cortex (dlPFC) and orbitofrontal cortex (OFC) is associated with psychopathic traits in a male criminal sample, suggesting poor performance in risky decision making and inhibition control for individuals high on psychopathic traits (Korponay et al., 2017). Moreover, psychopathic traits are associated with dysfunction of the limbic system, including the cingulate cortex, insula, and parahippocampal regions (Kiehl, 2006). A recent study reported decreased RSFC between the prefrontal and limbic-paralimbic regions for individuals with elevated psychopathic traits in a criminal sample, resulting in a weaker emotional and cognitive integration (Contreras-Rodríguez et al., 2015).

Furthermore, abnormal functional connectivity within and between functional brain networks has been reported to be associated with psychopathic traits (Espinoza et al., 2018; Thijssen & Kiehl, 2017). The functional brain networks refer to a collection of interconnected brain regions that interact to perform circumscribed functions (Bressler & Menon, 2010) and are related to a broad range of cognitive and affective functions (Raz, 2004; Sheline et al., 2009). Further, previous studies have indicated that the default mode network (DMN), executive-control /frontoparietal network (FPN), and salience/cingulo-opercular network (CON) may be particularly associated with psychopathic traits (Cohn et al., 2015; C. L. Philippi et al., 2015). For example, an neuroimaging study reported associations between psychopathic traits and power spectra (reflecting the degree of fluctuation in amplitudes of the intrinsic activity within the network) of the DMN and spatial maps (resembling the correspondence between a voxel time course and a region time course) of the FPN and CON in incarcerated adolescents, implicating deficits in self-referential processing, social cognition, and emotion regulation for adolescents with elevated psychopathic traits (Thijssen & Kiehl, 2017).

Although a plethora of evidence has identified abnormal neural correlates of extremely high psychopathic traits by utilizing RSFC in the correctional samples which only represent a small part in the psychopathy spectrum (Johanson, Vaurio, Tiihonen, & Lähteenvuo, 2020), it remains unclear whether these findings can be generalized to the general population with elevated psychopathic traits. Moreover, quantitative prediction of psychopathic traits at the individual level has not yet been investigated. Compared to the group-level analyses, predictions at the individual level of psychopathic traits could provide better clinical implications in the detection and potential intervention of psychopathic tendency (Lueken et al., 2016).

Connectome-based predictive modeling (CPM) —a data-driven machine learning approach— has been widely applied in predicting human behavior and personality traits by using whole-brain RSFC data (Finn et al., 2015; Shen et al., 2017). Compared with conventional correlation or regression analyses, CPM sufficiently extracts brain connectivity data to build predictive models and employs cross-validation to protect against overfitting (Shen et al., 2017; Yip, Scheinost, Potenza, & Carroll, 2019). This method has been well-validated in generating predictive model based on neural features (Feng, Wang, Li, & Xu, 2019; Ren et al., 2020). For example, CPM has been used to predict temperament (Jiang et al., 2018) and propensity to trust (Lu et al., 2019) based on whole-brain RSFC.

In this study, we employed CPM to predict individual differences of psychopathic traits measured by the Levenson Self-Report Psychopathy Scale (LSRP) based on whole-brain RSFC in a college student sample. Based on previous findings, we hypothesized that the connectome consisting of decreased RSFC involving the prefrontal cortex and limbic system is able to predict psychopathic traits at the individual level (N. E. Anderson & Kiehl, 2012; Thijssen & Kiehl, 2017). Further, given distinctions between primary and secondary psychopathy in behavioral and psychological features, we expected that the connectome in predicting primary psychopathy might be constituted by RSFC mainly involving regions (e.g., amygdala, insula and PCC) related to emotional function. However, the connectome in predicting secondary psychopathy might be constituted by RSFC mainly involving regions (e.g., OFC, dlPFC, and ACC) related to executive control. In addition, based on limited evidence on the association between psychopathic traits and large-scale networks, we expected RSFC within and between the three large-scale networks—FPN, DMN and CON to be able to predict individual psychopathic traits.

## Method

### Participants

Eighty-four healthy right-handed college students (mean age: 22.6 ± 2.59 years, 34 males and 50 females) were recruited from Hangzhou Normal University in China. All participants had no history of neurological or psychiatric disorders. This study was conducted in accordance with the Helsinki Declaration. Written informed consent was obtained from each participant for this study, which was approved by the Ethics Committee of Hangzhou Normal University.

### Measures

Psychopathic traits were assessed using the LSRP (Levenson, Kiehl, & Fitzpatrick, 1995). The LSRP is a 26-item rating scale with two subscales assessing primary and secondary psychopathy, and each of the items can be rated on a 4-point Likert agree/disagree scale (1 [strongly disagree] to 4 [strongly agree]). The total LSRP score represents the overall level of psychopathic traits (i.e., total psychopathic trait score, T-PTS). The primary psychopathy subscale assesses the *affective traits of psychopathy* such as lack of empathy and shallow affect (i.e., primary psychopathic trait score, P-PTS). The secondary psychopathy subscale assesses the *antisocial traits of psychopathy*, such as impulsiveness and antisocial behavior (i.e., secondary psychiatric trait score, S-PTS) (Miller, Gaughan, & Pryor, 2008). The LSRP has been validated in Chinese university student samples (Wang et al., 2018). The Cronbach’s Alphas were 0.80 (T-PTS), 0.79 (P-PTS), and 0.59 (S-PTS) in the present sample.

### Image data acquisition

Imaging data were collected at the Center for Cognition and Brain Disorders of Hangzhou Normal University using a 3T MRI scanner (GE Discovery 750 MRI, General Electric, Milwaukee, WI, USA). Participants were instructed to keep their eyes closed and stay awake without performing any cognitive exercises during the 8-minute RS scan. The imaging parameters of the EPI sequence including 240 images were as follows: repetition time (TR) = 2000 ms; interleaved 43 slices; echo time (TE) = 30 ms; thickness = 3.2 mm; flip angle = 90°; field of view (FOV) = 220 ×220 mm^2^; and matrix size = 64 × 64. A high-resolution 3D T1-weighted anatomical image was also acquired through magnetization-prepared rapid acquisition with gradient-echo (MPRAGE) sequence with following parameters: 176 sagittal slices; TR = 8100 ms; slice thickness = 1 mm; TE = 3.1 ms; flip angle = 8°; FOV = 250 × 250 mm. This session lasted for about 5◻minutes.

### Preprocessing

The fMRI data preprocessing was performed with DPABI v3.0 (Data processing & Analysis for Brain Imaging, http://rfmri.org/dpabi) (**Fig.1a**) (Yan, Wang, Zuo, & Zang, 2016). The first 10 volumes were discarded for signal equilibrium and participants’ adaptation to the scanning environment. The remaining 230 images were subsequently realigned for head movement correction and spatially normalized into the standard Montreal Neurological Institute space (MNI template, resampling voxel size was 3◻×◻3◻×◻3 mm^3^). Next, the linear trends of time courses were removed, and a band◻pass temporal filtering (0.01–0.1 Hz) was applied to the time series to reduce the effect of low◻frequency drifts and high◻frequency physiological noise. Finally, the images were smoothed using a Gaussian filter with a kernel of full width at half maximum of 4 mm, and three common nuisance variables (i.e., white matter signal, cerebrospinal fluid signal, and 24 movement regressors including 6 standard head motion parameters, the derivative of the standard motion parameters, and the 12 corresponding squared items) were regressed out.

### RSFC

Whole-brain RSFC was computed by employing the Dosenbach atlas with 142 nodes excluding the cerebellum (**Supplementary Table 1** & **Fig. 1b**) (Dosenbach et al., 2010). Every node defined in the atlas was a 5 mm-radius spherical region can be assigned into five pre-defined networks: default-mode (DMN), frontoparietal (FPN), cingulo-opercular (CON), sensorimotor (SMN), and occipital (OCCN) network. We extracted the time courses of all nodes by averaging the time course for all the voxels within a node. Pearson’s correlation coefficient between the time courses of each pair of nodes was calculated and subsequently was transformed using Fisher’s z-transformation. A symmetric 142×142 connectivity matrix was constructed for each subject in which each element represents RSFC strength between two nodes.

**Figure 1.**
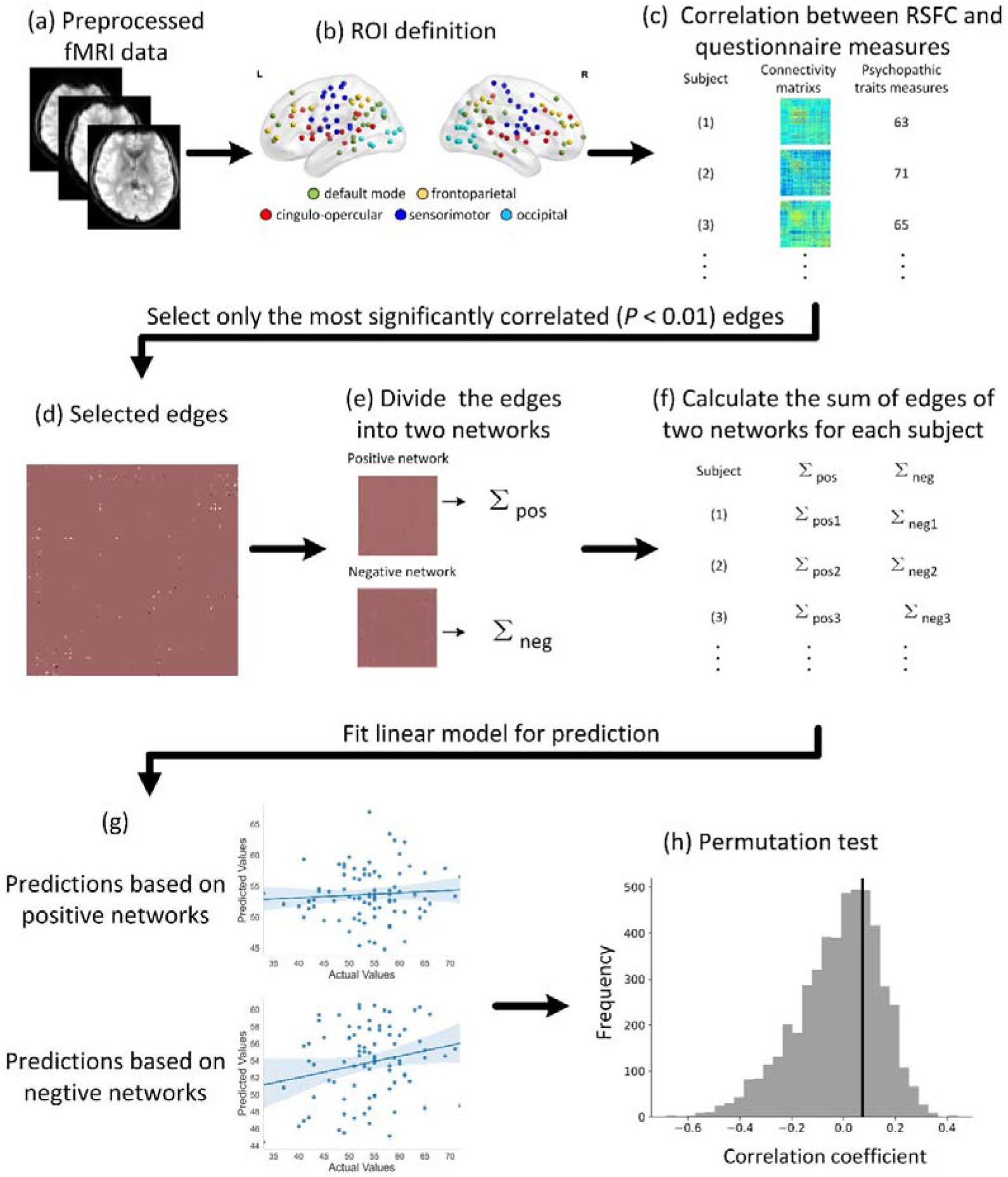
The workflow of connectome-based predictive modeling (CPM). Preprocessed resting-state fMRI data (a) were used to calculate whole-brain functional connectivity based on the Dosenbach atlas (b). Correlation analyses were then performed between connectivity matrix and questionnaire measurements (e.g., total LSRP score) (c). The important edges were selected by setting a threshold of *p*<0.01(d). The edges were separated into the positive network (edges positively correlated with the questionnaire measurements) and negative network (edges negatively correlated with behavioral measurements) (e). For each participant, the sum of selected edges for the two networks were calculated (f). Linear predictive models were built to predict psychopathic traits. The CPM made out-of-sample predictions by applying a cross-validated approach (g). A permutation test was utilized to examine the performance of predictive models (h).

### CPM

All the analyses listed below used the CPM framework to construct predictive models predicting the targets (S-PTS, S-PTSS, S-PTS) based on whole brain RSFC using MATLAB (http://www.mathworks.com) scripts taken from Shen’s paper published in 2017 (https://www.nitrc.org/projects/bioimagesuite). First, Pearson’s correlation coefficients between each edge and targets were calculated across the participants (**Fig. 1c**). Edges in which the p-values of the correlation were less than the threshold (*p* < 0.01) were defined as important edges for the subsequent processing (**Fig. 1d**). The positive (negative) networks were composed of edges that were positively (negatively) associated with psychopathic traits (**Fig. 1e**). Next, the edge weights (connectivity strength) were summed up in positive and negative networks respectively for each participant, ultimately forming single-subject positive and negative network strengths (**Fig. 1f**). Then, linear regression predictive models were constructed to predict psychopathic traits by including positive and negative network strengths as regressors. Finally, the resultant linear equation including slope and intercept was applied in new participants to obtain predictive values (**Fig. 1g**).

### Cross-validation

A leave-one-subject-out cross-validation (LOOCV) approach was utilized to test the generalizability of the prediction. N-1 participants were set as the training set (N equals to the number of the participants) while the left-out participants were set as the testing set. The predictive model was built using the training data and then was applied to predict the testing participants’ behavior. The training and testing procedures were implemented in an iteration manner until all the participants have a predictive value.

Moreover, model performance was evaluated with Pearson’s correlation coefficient (*r*) and standard mean square error (*SMSE*) between the predictive value and actual value while controlling for confounding variables such as age, gender, and head motion. The significance of *r* and *SMSE* was assessed with a permutation test of 5,000 permutations. Null distributions were generated by randomly shuffling the actual target value 5,000 times and repeating the CPM analyses with the shuffled data. The number of permutations with better performance (i.e., higher *r* and lower *SMSE*) than the model built with the true actual value were calculated and used to divide the total number of permutations for obtaining the *p*-value.

### Node and network analyses

After the cross-validation, consensus functional connectivity that retained in every iteration was obtained, and the degree centrality (i.e., the number of connections involving a certain node) of nodes involved in consensus functional connectivity was calculated. The first eight nodes with the largest degree centrality were defined as the most connected nodes in the predictive connectivity. Also, the connections between and within five large-scale brain networks (i.e., CON, DMN, FPN, OCCN, and SMN) were computed to characterize the predictive connectome at the network level, and the inter- or intra-network connections with the largest number regarded as connections that most contribute to the predictive connectome.

### Validation analyses

To avoid the overfitting of the predictive model, k-fold cross-validations —2-, 5-, and 10-fold cross-validations— were further applied to validate the results (Poldrack, Huckins, & Varoquaux, 2020). Like LOOCV, k-fold cross-validation divides the subjects randomly and equally into k subsets in which one subset was used as the testing set, and other remaining subsets were used as the training set. The predictive model was trained by the training set and was then used to make predictions for the testing set. The procedure repeated k times to ensure each subset was used as the testing set once. Pearson correlation coefficients between the predictive and actual value were calculated to evaluate the model performance. Given that all the subjects were randomly divided into k subsets and model performance is largely affected by the data division, the k-fold cross-validation was repeated 50 times to increase reliability. The correlation coefficients were averaged to obtain a final prediction performance, and a permutation test was utilized 5,000 times to test the significance of the model performance.

## Results

Psychopathic traits (i.e., T-PTS, P-PTS, S-PTS) were not significantly correlated with age (T-PTS : *r* = −0.033, *p* = 0.764; P-PTS: *r* = 0.027, *p* = 0.807; S-PTS : *r* = −0.116, *p* = 0.295) and no significant gender differences were found (T-PTS: *t* = 0.633, *df* = 82, *p* = 0.529; P-PTS: *t* = −0. 153, *df* = 82, *p* = 0.879; S-PTS: *t* = 1.657, *df* = 82, *p* = 0.101) (**Tab. 1**).

**Table 1.** Participants demographics and LSRP scores.

|  | Male (N=34) |  | Female (N=50) |  | *p*-value |
| --- | --- | --- | --- | --- | --- |
|  | Mean | SD | Mean | SD |  |
| Age (years) | 23.29 | 3.13 | 22.10 | 2.05 | 0.056 |
| T-PTS | 53.15 | 8.02 | 54.22 | 7.35 | 0.529 |
| P-PTS | 33.27 | 5.93 | 33.08 | 5.07 | 0.879 |
| S-PTS | 19.88 | 3.21 | 21.14 | 3.55 | 0.101 |

SD: standard deviation; T-PTS: total psychopathic trait score; P-PTS: primary psychopathic trait score; S-PTS: secondary psychopathic trait score.

The CPM results revealed that the negative network strength significantly predicted T-PTS (*r* = 0.295, *p(r)* = 0.011, *SMSE* = 1.485, *p(SMSE)* = 0.022, **Fig. 2a**) and S-PTS (*r* = 0.309, *p* = 0.008, *SMSE* = 0.773, *p(SMSE)* = 0.014, **Fig. 2c**) but not P-PTS (*r* = 0.062, *p* = 0.309, *SMSE* = 1.234, *p(SMSE)* = 0.370, **Fig. 2b**) after controlling for the potential confounding effect of gender, age and head motion. In contrast, the positive network strength could not reliably predict any of the targets (T-PTS : *r* = 0.073, *p(r)* = 0.308, *SMSE* = 1.771, *p(SMSE)* = 0.353; S-PTS : *r* = 0.129, *p* = 0.199, *SMSE* = 0.903, *p(SMSE)* = 0.233; P-PTS: *r* = 0.008, *p* = 0.468, *SMSE* = 1.214, *p(SMSE)* = 0.425). Across all folds of cross-validation, 78 edges (4 positive and 74 negative), 34 edges (3 positive and 31 negative), and 98 edges (3 positive and 95 negative) retained as consensus functional connectivity for predicting T-PTS, P-PTS, and S-PTS, respectively. Because only a few edges in the positive tail retained in all the folds of cross-validation and could not provide reliable predictions for any of the targets, the following analyses were only performed only for the negative networks.

**Figure 2.**
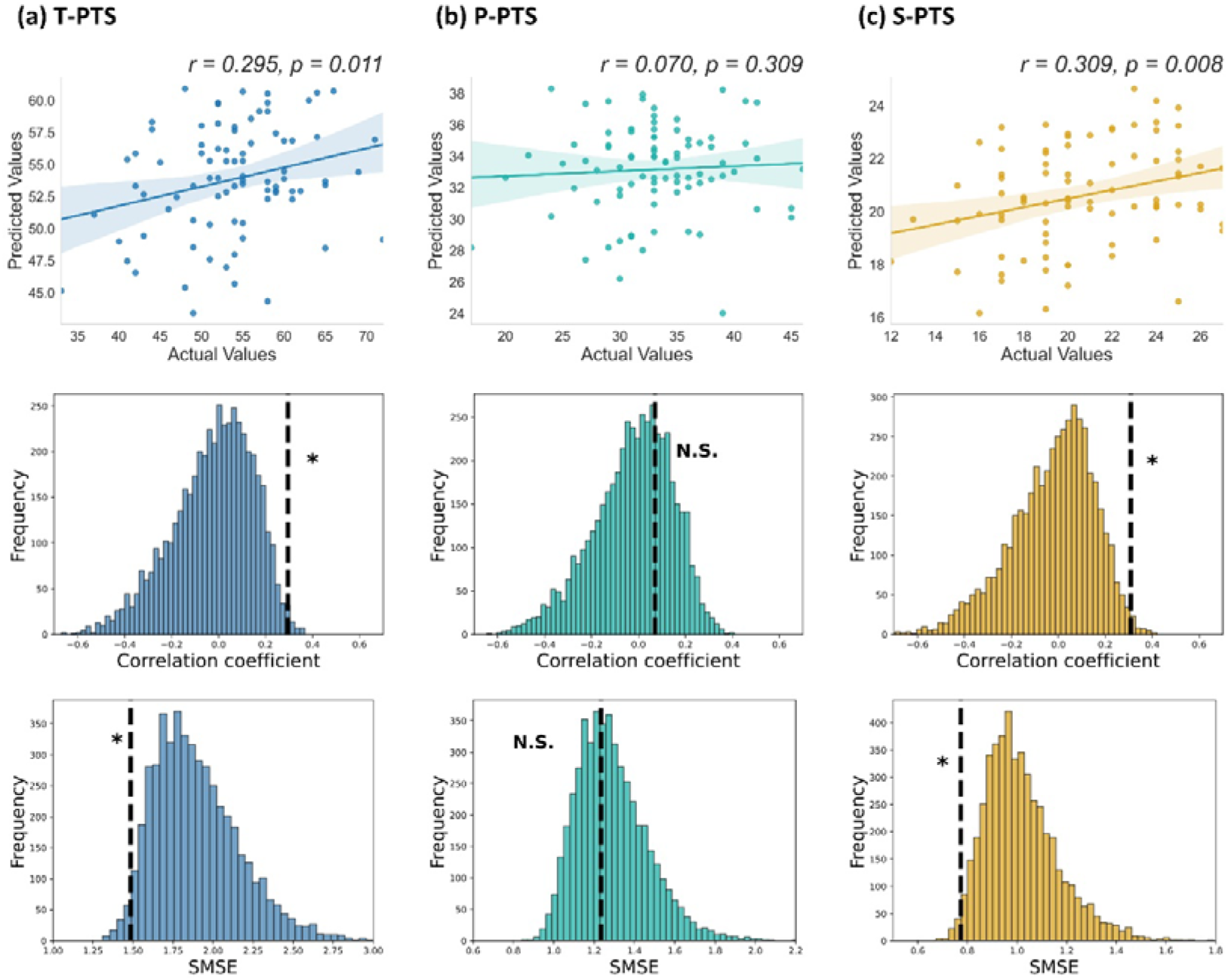
Performance of the prediction models for total LSRP, primary psychopathy, and secondary psychopathy score based on the negative network. Correlation between actual and predicted values, and permutation distribution of the correlation coefficient and SMSE for the prediction of the total psychopathic traits score (a), primary psychopathic traits score (b), and secondary psychopathic traits score (c). The values obtained using the real scores are indicated by the black dash line. T-PTS: total psychopathic trait score; P-PTS: primary psychopathic trait score; S-PTS: secondary psychopathic trait score; SMSE: standard mean square error; N.S.: not significant; * *p* < 0.05.

Figure 3 summarizes the negative networks predicting T-PTS and S-PTS. For T-PTS, the most highly connected nodes included the superior temporal sulcus (STS), middle occipital gyrus (MOG), anterior prefrontal cortex (aPFC), inferior orbitofrontal cortex (IOFC), vmPFC, insula, and supramarginal gyrus (SMG) (**Tab. 2**). For S-PTS, the most highly connected nodes included the aPFC, superior frontal gyrus (SFG), MOG, OFC, calcarine, and anterior cingulate cortex (ACC) (**Tab. 3**). In terms of large-scale functional networks, the connections of the CON with OCCN predicted T-PTS and S-PTS (**Fig. 4**).

**Table.2.**
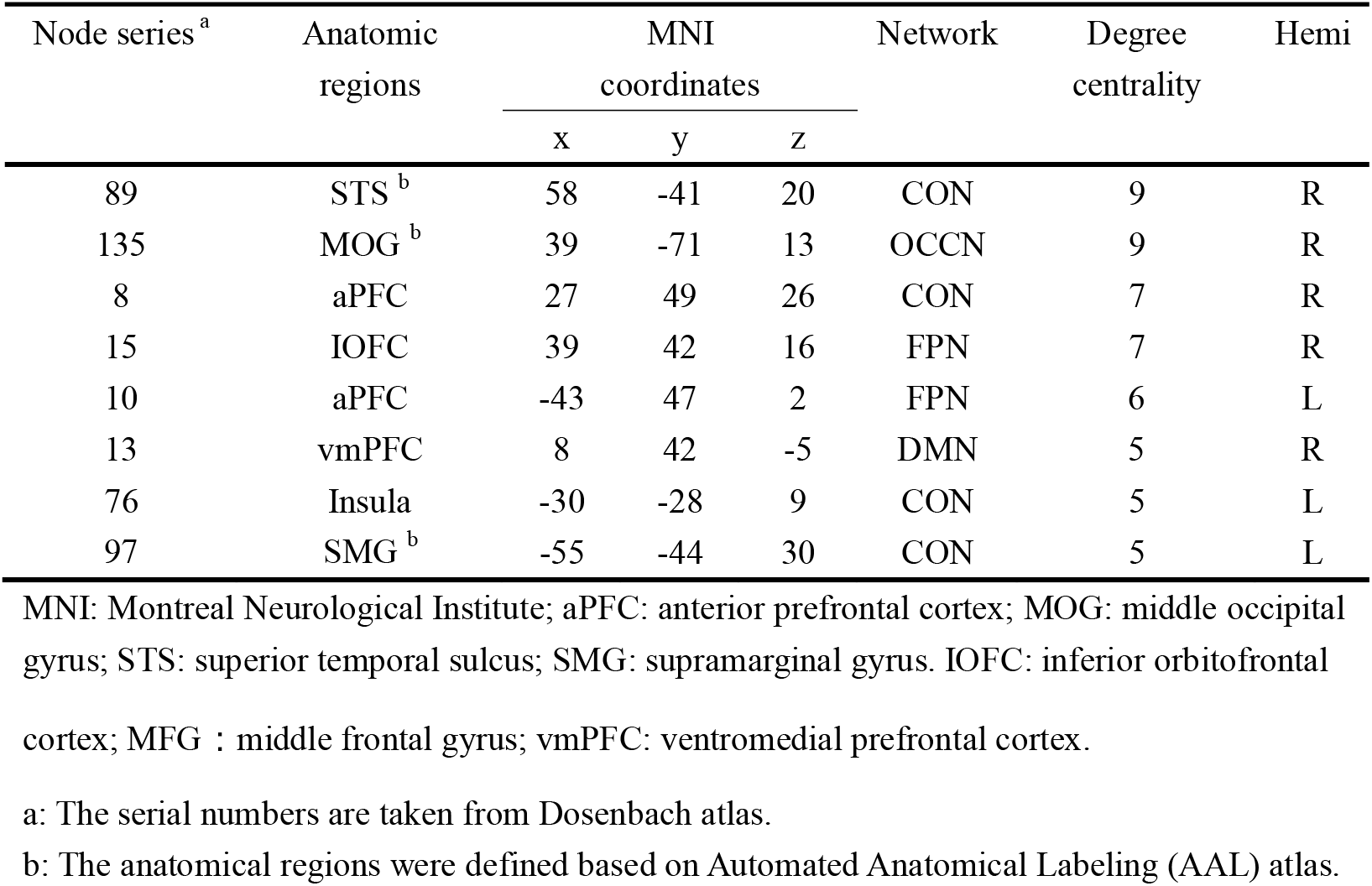
Eight nodes (5%) with the most connections in the negative networks in predicting total psychopathic trait score.

| Node series <sup>a</sup> | Anatomic regions | MNI coordinates |  |  | Network | Degree centrality | Hemi |
| --- | --- | --- | --- | --- | --- | --- | --- |
|  |  | x | y | z |  |  |  |
| 89 | STS <sup>b</sup> | 58 | -41 | 20 | CON | 9 | R |
| 135 | MOG <sup>b</sup> | 39 | -71 | 13 | OCCN | 9 | R |
| 8 | aPFC | 27 | 49 | 26 | CON | 7 | R |
| 15 | IOFC | 39 | 42 | 16 | FPN | 7 | R |
| 10 | aPFC | -43 | 47 | 2 | FPN | 6 | L |
| 13 | vmPFC | 8 | 42 | -5 | DMN | 5 | R |
| 76 | Insula | -30 | -28 | 9 | CON | 5 | L |
| 97 | SMG <sup>b</sup> | -55 | -44 | 30 | CON | 5 | L |

MNI: Montreal Neurological Institute; aPFC: anterior prefrontal cortex; MOG: middle occipital gyrus; STS: superior temporal sulcus; SMG: supramarginal gyrus. IOFC: inferior orbitofrontal cortex; MFG : middle frontal gyrus; vmPFC: ventromedial prefrontal cortex.
a: The serial numbers are taken from Dosenbach atlas.
b: The anatomical regions were defined based on Automated Anatomical Labeling (AAL) atlas.

**Table.3.** Eight nodes with the most connections in the negative networks in predicting secondary psychopathic trait score.

| Node series <sup>a</sup> | Anatomic regions | MNI<br>coordinates |  |  | Network | Degree<br>centrality | Hemi |
| --- | --- | --- | --- | --- | --- | --- | --- |
|  |  | x | y | z |  |  |  |
| 8 | aPFC | 27 | 49 | 26 | CON | 12 | R |
| 10 | aPFC | -43 | 47 | 2 | FPN | 12 | L |
| 50 | SFG <sup>b</sup> | -26 | -8 | 54 | SMN | 10 | L |
| 135 | MOG <sup>b</sup> | 39 | -71 | 13 | OCCN | 8 | R |
| 9 | OFC <sup>b</sup> | 42 | 48 | -3 | FPN | 8 | R |
| 152 | Calcarine <sup>b</sup> | -5 | -80 | 9 | OCCN | 7 | L |
| 21 | ACC | -1 | 28 | 40 | FPN | 7 | L |
| 19 | ACC | -2 | 30 | 27 | CON | 6 | L |

MNI: Montreal Neurological Institute; aPFC: anterior prefrontal cortex; SFG: superior frontal gyrus; MOG: middle occipital gyrus; OFC: orbitofrontal cortex; ACC: anterior cingulate cortex.
a: The serial numbers are taken from Dosenbach's atlas.
b: The anatomical regions were defined based on Automated Anatomical Labeling (AAL) atlas.

**Figure 3.**
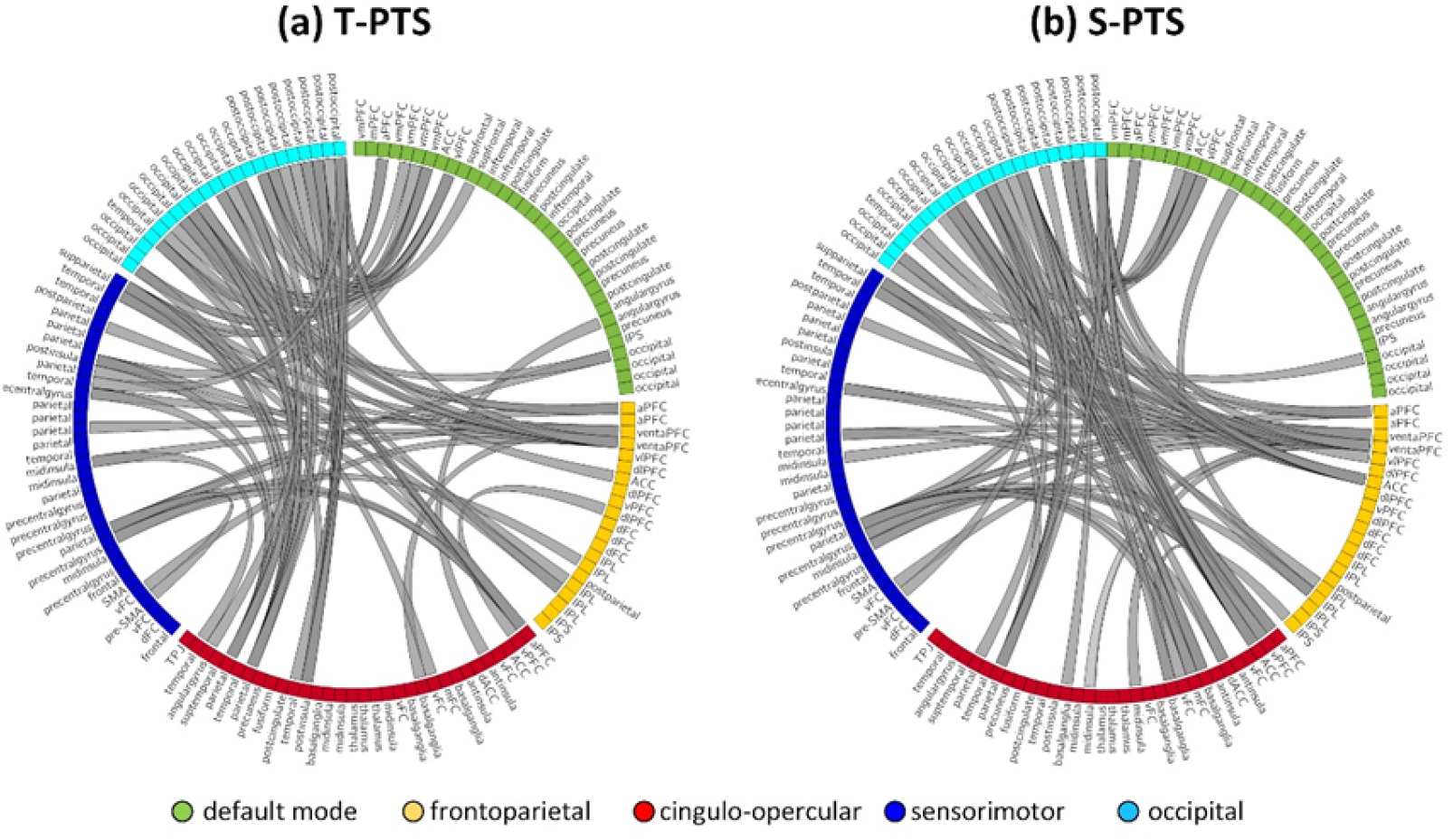
Negative networks for predicting total LSRP and secondary psychopathy score. Negative networks in predicting total psychopathic trait score (a) and secondary psychopathic trait score (b). T-PTS: total psychopathic trait score; S-PTS: secondary psychopathic trait score.

**Figure 4.**
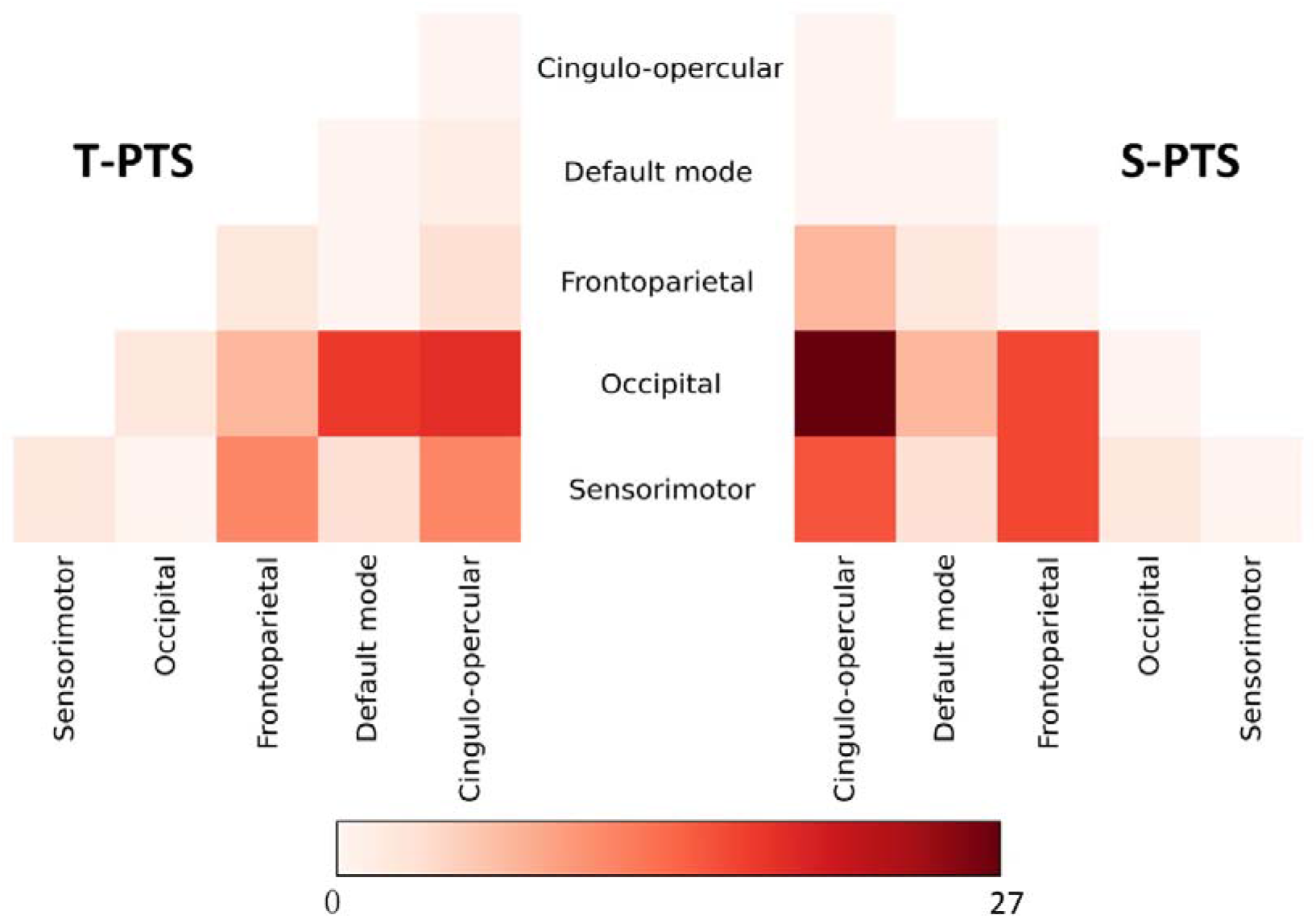
Within- and between-network connectivity for the negative networks. Cells represent the total number of edges connecting nodes within (and between) each network, with darker colors indicating a greater number of edges. T-PTS: total psychopathic trait score; S-PTS: secondary psychopathic trait score.

The predictive model performance was further validated and remained significant using the k-fold cross-validations (i.e., 2-, 5-, and 10-fold cross-validations) (**Tab. 4**).

**Table 4.**
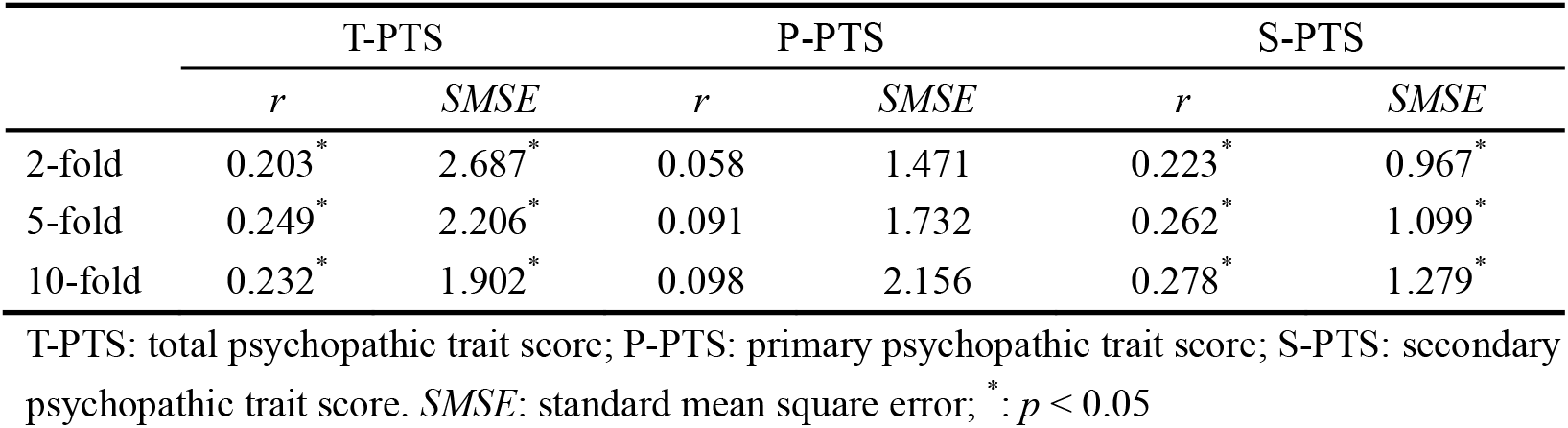
Results of validation analyses.

|  | T-PTS |  | P-PTS |  | S-PTS |  |
| --- | --- | --- | --- | --- | --- | --- |
|  | *r* | *SMSE* | *r* | *SMSE* | *r* | *SMSE* |
| 2-fold | 0.203<sup>*</sup> | 2.687<sup>*</sup> | 0.058 | 1.471 | 0.223<sup>*</sup> | 0.967<sup>*</sup> |
| 5-fold | 0.249<sup>*</sup> | 2.206<sup>*</sup> | 0.091 | 1.732 | 0.262<sup>*</sup> | 1.099<sup>*</sup> |
| 10-fold | 0.232<sup>*</sup> | 1.902<sup>*</sup> | 0.098 | 2.156 | 0.278<sup>*</sup> | 1.279<sup>*</sup> |

T-PTS: total psychopathic trait score; P-PTS: primary psychopathic trait score; S-PTS: secondary psychopathic trait score. *SMSE*: standard mean square error; <sup>\*</sup>: $p < 0.05$

## Discussion

In the present study, we applied CPM to predict individual variations in psychopathic traits (i.e., T-PTS, P-PTS, S-PTS). We demonstrated that whole-brain RSFC connectome predicted T-PTS and S-PTS but not P-PTS at the individual level. Inter-individual variations in psychopathic traits were predicted by RSFC connectome. Particularly, the most connected nodes located in the PFC (i.e., aPFC, OFC, and vmPFC) and limbic system (e.g., insula and ACC). In addition, predictive connectomes were primarily constituted by connections between OCCN and CON at the level of large-scale functional networks.

Psychopathic traits have been recognized to increase the risk of violence and aggression (Porter, Woodworth, & Black, 2018). Individuals with elevated psychopathic traits exhibit deficient cognitive, affective, and social functions which may be related to their abnormal neurobiological correlates (Deming & Koenigs, 2020). Here, we demonstrated that CPM based on the whole-brain RSFC successfully predicted T-PTS and S-PTS at the individual level. However, only the negative networks in the predictive models made significant predictions whereas the positive networks failed to successfully predict psychopathic traits due to the limited number of edges. Our findings added to previous evidence that decreased RSFC involving brain regions related to emotional, social, and cognitive function (Contreras-Rodríguez et al., 2015; Espinoza et al., 2019), indicating that higher psychopathic traits may be primarily accompanied by the decline of the intrinsic whole-brain RSFC which might contribute to functional segregation of brain regions.

However, the whole-brain RSFC failed to predict P-PTS which has been typically associated with regions supporting emotional function such as amygdala and vmPFC. For example, attenuation of amygdala reactivity during emotion recognition and evocation were observed in psychopaths (Gordon, Baird, & End, 2004; Marsh & Cardinale, 2014). A rs-fMRI study found decrease in FC between the amygdala and the vmPFC in individuals with psychopathy (Motzkin et al., 2011). In particular, weakened amygdala-vmPFC FC contributes to psychopathic symptoms in low-income males (Waller et al., 2019). Here, the whole-brain RSFC was not able to predict primary, which is likely due to that the college student sample with elevated psychopathic traits in this study may not have emotional deficits as severe as clinical psychopaths.

On the one hand, we found that the high-degree nodes in the T-PTS predictive model were primarily anchored in the aPFC, OFC, vmPFC which have been consistently associated with psychopathic traits. The aPFC is frequently engaged in cognitive processing including goal-oriented planning (Koechlin, Basso, Pietrini, Panzer, & Grafman, 1999), reallocation of attention (Ramnani & Owen, 2004), and impulse control (Ramnani & Owen, 2004). The OFC is responsible for inhibitory control, learning and value encoding (Rudebeck & Rich, 2018) and plays an important role in regulating emotional responses such as fear and anxiety since it strongly connects with amygdala and insula (Rudebeck, Saunders, Prescott, Chau, & Murray, 2013). The vmPFC is involved in the reinforcement-based decision making which is crucial for norm compliance (R. J. R. Blair, 2007). Dysfunction of the vmPFC contributes to immoral behavior that is commonly seen in psychopathic individuals (R. Blair, 2011; Koenigs, Kruepke, Zeier, & Newman, 2012). In general, abnormal interactions of the regions in the prefrontal cortex may contribute to poor self-control, impulsivity and aggressivity associated with psychopathic traits (Hoppenbrouwers et al., 2013).

On the other hand, we demonstrated that high-degree nodes in the negative networks involved regions supporting emotional and social processing, such as the insula, the STS and SMG. The insula has been widely recognized to be involved in emotional processing (Menon & Uddin, 2010). Activations of the insula during an emotional facial expression task are negatively correlated with psychopathic traits (Seara-Cardoso, Sebastian, Viding, & Roiser, 2016). Our results indicate that altered intrinsic network architecture involving the insula may be related to atypical emotional functions for individuals with higher psychopathic traits. The STS and SMG is engaged in social cognition (Zilbovicius et al., 2006) and especially plays an important role in overcoming emotional egocentricity (Isik, Koldewyn, Beeler, & Kanwisher, 2017; Silani, Lamm, Ruff, & Singer, 2013). Dysfunction of the connection involving these two regions may underlie the abnormal social processing of individuals with elevated psychopathic traits.

Secondary psychopathy has been suggested to be associated with impulsivity and antisocial behavior (Dean et al., 2013). Similar to the T-PTS predictive model, nodes located in the prefrontal cortex (i.e., aPFC) also had a relatively high degree in the S-PTS predictive model. Additionally, the ACC served as another high-degree node in the model that significantly predicted secondary psychopathy. The ACC is implicated in the emotional regulation (Hornak et al., 2003), reward assessment (Knutson, Westdorp, Kaiser, & Hommer, 2000), and risky decision-making (Fishbein et al., 2005). Accumulating evidence indicated that the ACC is robustly related to impulse control which is involved in multiple processes including response inhibition and affective decision making (Castellanos-Ryan & Séguin, 2016; Hoerst et al., 2010; Strafella, 2019). Thus, dysfunction of the ACC may underlie insufficient decision-making and poor impulsive control in individuals with high secondary psychopathy score (Koenigs, 2012).

Additionally, it is worth noting that connections between the OCCN and CON constituted the predictive models of T-PTS and S-PTS in terms of the large-scale functional networks. The OCCN is typically involved in visual functions such as visual perception and discrimination (Likova & Tyler, 2008), and the CON is mostly engaged in saliency detention and emotional processing (Coste & Kleinschmidt, 2016; Sadaghiani & D’Esposito, 2015). The amygdala, a key node in the CON has extensive connections with visual regions and the functional connections between OCCN and CON may reflect the emotion-related modulation of visual processing stream (Morris, Buchel, & Dolan, 2001). It has been previously suggested that psychopathy scores were associated with abnormal connectivity within and between the FPN, DMN and CON (Carissa L Philippi et al., 2015; Thijssen & Kiehl, 2017).

Here, we provided novel evidence that decreased OCCN-CON intrinsic functional connectivity was associated with higher psychopathic traits, reflecting atypical visual attentional processing to emotionally salient stimuli in the environment for individuals with elevated psychopathic traits. Consistent with this assumption, psychopathic traits have been shown to be associated with reduced augmentation effect of emotion content on the brain activity in visual processing regions in a task which requires attention to emotional features of visual stimuli, suggesting abnormal interaction between visual attention and emotional processing for individuals with elevated psychopathic traits (Nathaniel E Anderson et al., 2017).

Several limitations in the current study should be addressed in future studies. First, the sample of the present study was college students with a relatively modest sample size (n=84). The college student sample is different from the community sample especially in age, social status, and intelligence level. Therefore, future work using a large community sample is warranted. Second, although the present study significantly predicted the T-PTS and S-PTS based on RSFC, the reliability of the results needs further validation. Future studies could use external samples to test the generalizability and validate the predictive models. Third, while we found that decreased OCCN-CON connections predicted psychopathic traits at the large-scale network level, it needs more evidence to verify the results as the OCCN has not been typically found to be associated with psychopathic traits. Despite these limitations, our study sheds new light on neural signatures of psychopathic traits in college students.

## Conclusion

The current study demonstrated that the connectome consisting of whole-brain RSFC successfully predicted T-PTS and S-PTS at the individual level in a college student sample. Notably, high-degree nodes in the predictive models mainly included brain regions located in the prefrontal cortex and limbic system. In addition, the OCCN-CON connections also constituted the predictive connectomes. The CPM approach provides a novel tool to characterize the neural underpinnings of psychopathic traits. Moreover, our findings may advance our understanding of psychopathic traits in the non-clinical sample and may have potential applications in the early diagnosis and intervention of psychopathy.

## Notes

### Competing Interest Statement

The authors have declared no competing interest.

